# Activation of innate immune signalling during development predisposes to inflammatory intestine and shortened lifespan

**DOI:** 10.1101/2021.04.12.439419

**Authors:** Kyoko Yamashita, Ayano Oi, Hina Kosakamoto, Toshitaka Yamauchi, Hibiki Kadoguchi, Takayuki Kuraishi, Masayuki Miura, Fumiaki Obata

## Abstract

Early-life inflammatory response is associated with risks of age-related pathologies. How transient immune signalling activity during animal development influences life-long fitness is not well understood. Using *Drosophila* as a model, we find that activation of innate immune pathway IMD signalling in the developing larvae increases adult starvation resistance, decreases food intake, and shortens organismal lifespan. Interestingly, lifespan is shortened by the IMD activation in the larval gut and fat body, while starvation resistance and food intake are altered by that in neurons. The adult flies developed with IMD activation show sustained IMD activity in the gut, despite complete tissue renewal during metamorphosis. The inflammatory adult gut is associated with a greater amount of *Gluconobacter* sp., characteristic gut microbiota increased in response to immune activation. Removing gut microbiota by antibiotics attenuates the increase of IMD activity and rescues the shortened lifespan. This study demonstrates a tissue-specific programming effect of early-life immune activation on the adult physiology and organismal lifespan.

## Introduction

Immunity needs to be tightly controlled as both shortages and excesses of immune activation are detrimental to organisms. Chronic, and often systemic, inflammatory response occurs during ageing, which can increase the risk of various age-related diseases [1]. *Drosophila melanogaster* is a genetically tractable model for studying how immune pathways are involved in the ageing process. The immune deficiency (IMD) pathway is an evolutionally conserved immune regulator in *Drosophila*, which is a counterpart of the tumour necrosis factor receptor (TNFR) pathway in mammals [2]. The IMD pathway is activated upon infection with bacteria possessing DAP-type peptidoglycan, and is known to be spontaneously activated in aged animals, at least partly in a gut microbiota-dependent manner. Removing microbiota or overexpressing negative regulators for the IMD pathway in the adult gut attenuates age-related IMD activation and concomitantly extends lifespan [3–5]. Activation of the IMD pathway in gut progenitor cells induces hyperproliferation of intestinal stem cells [6]. A chronic inflammatory condition in aged flies triggers neurodegeneration and shortens lifespan, which can be rescued by inhibiting IMD signalling in glia cells [7]. Age-related activation of the IMD pathway in the renal (Malpighian) tubules induced by a commensal *Acetobacter persici* triggers age-dependent metabolic shifts, including purine metabolism [8]. These studies have revealed that age-related immune activation in various tissues leads to organismal ageing.

Early-life environmental stressors have prolonged effects on adult health, as often described as the “Developmental Origins of Health and Diseases (DOHaD) hypothesis” [9–11]. In *Drosophila*, dietary protein restriction only in the larval stage extends lifespan via altered lipid metabolism [12]. Developmental exposure to low-dose oxidant remodels the gut microbiome and extends lifespan [5]. Hypoxic conditions during development decrease starvation resistance and lifespan [13]. These studies illustrate how environmental factors during development can program adult physiology and lifespan.

Various stressors regarded as risk factors for age-related diseases, such as malnutrition, irradiation, chemical exposures, smoke, alcohol, or even mental stress, commonly lead to inflammatory response. A longitudinal cohort study suggested that childhood infection is correlated with the incidence of cardiovascular diseases in 40-year old humans [14]. This and many other epidemiological studies have implied that early-life inflammation is associated with inflammatory diseases and mortality in adulthood [15,16], however few studies directly test the causal relationship. Irradiation during development increases cell death in the adult brain and decreases locomotive ability and organismal lifespan in *Drosophila* [17]. In this condition, persistent immune activation is observed in adult flies [18]. On the other hand, oral infection of *Erwinia carotovora* (Ecc15) in larvae does not affect the lifespan of adult flies [19]. Genetic manipulation is also useful to test how early-life signalling activity impacts adult lifespan. For example, loss of function of *PGAM5* triggers immune-related genes in larvae, which is associated with increased lifespan through a prolonged FoxO activation [20]. Decreasing mitochondrial electron transport by knocking down *ND75* specifically in muscles on the first day of the larval stage can extend lifespan [21]. It is likely that infection as well as other stressors trigger not only immune activation but also complex reactions such as tissue injury/recovery response or metabolic remodelling. Thus, despite the implication that early-life immune response affects organismal lifespan, whether an immune signalling activity *per se* during development influences the lifespan and adult physiology has not been directly assessed.

In this study, we attempted to test whether immune activation in a larval stage-restricted manner can alter adult fitness and lifespan. We found, in adult flies with larval IMD activation, that immune and metabolic alteration occurs and shortens organismal lifespan.

## Results

### Establishment of mild immune activation during development

We used the Gene Switch (GS) system to achieve precise control of gene manipulation [22]. GS is useful to induce any gene of interest by an inducer RU486 in a dose-dependent manner. For activation of the innate immune IMD pathway in larvae, we overexpressed constitutive active form of IMD (*IMD^CA^*) using a ubiquitous driver *Daughterless GS* (*Da^GS^*). *IMD^CA^* lacks the N-terminal inhibitory domain and therefore is active in the absence of bacterial stimulation [6]. We put embryos of *Da^GS^>IMD^CA^* and its negative control *Da^GS^>LacZ* on top of the standard *Drosophila* diets containing various doses of RU486, and allowed them to develop into adult flies (Fig. 1A). Feeding 200 μM of RU486, the concentration frequently used for adult flies, caused higher developmental lethality even for the control flies in our laboratory condition. Decreasing the RU486 concentration to 5 μM resulted in little effect on viability and adult body weight for the control (*Da^GS^>LacZ*) animals, but strong lethality and decreased body weight for *Da^GS^>IMD^CA^*, suggesting that IMD activation impairs larval growth (Fig. 1B, C). When the concentration of RU486 was decreased to 1 μM, adult *Da^GS^>IMD^CA^* flies showed normal body weight (Fig. 1C). At this concentration, we observed mild developmental delay compared with the control (no RU486 treatment), but this was rather due to a side effect of RU486, as the phenotype was also obvious for *Da^GS^>LacZ* flies (Fig. S1A).

**Fig. 1.**
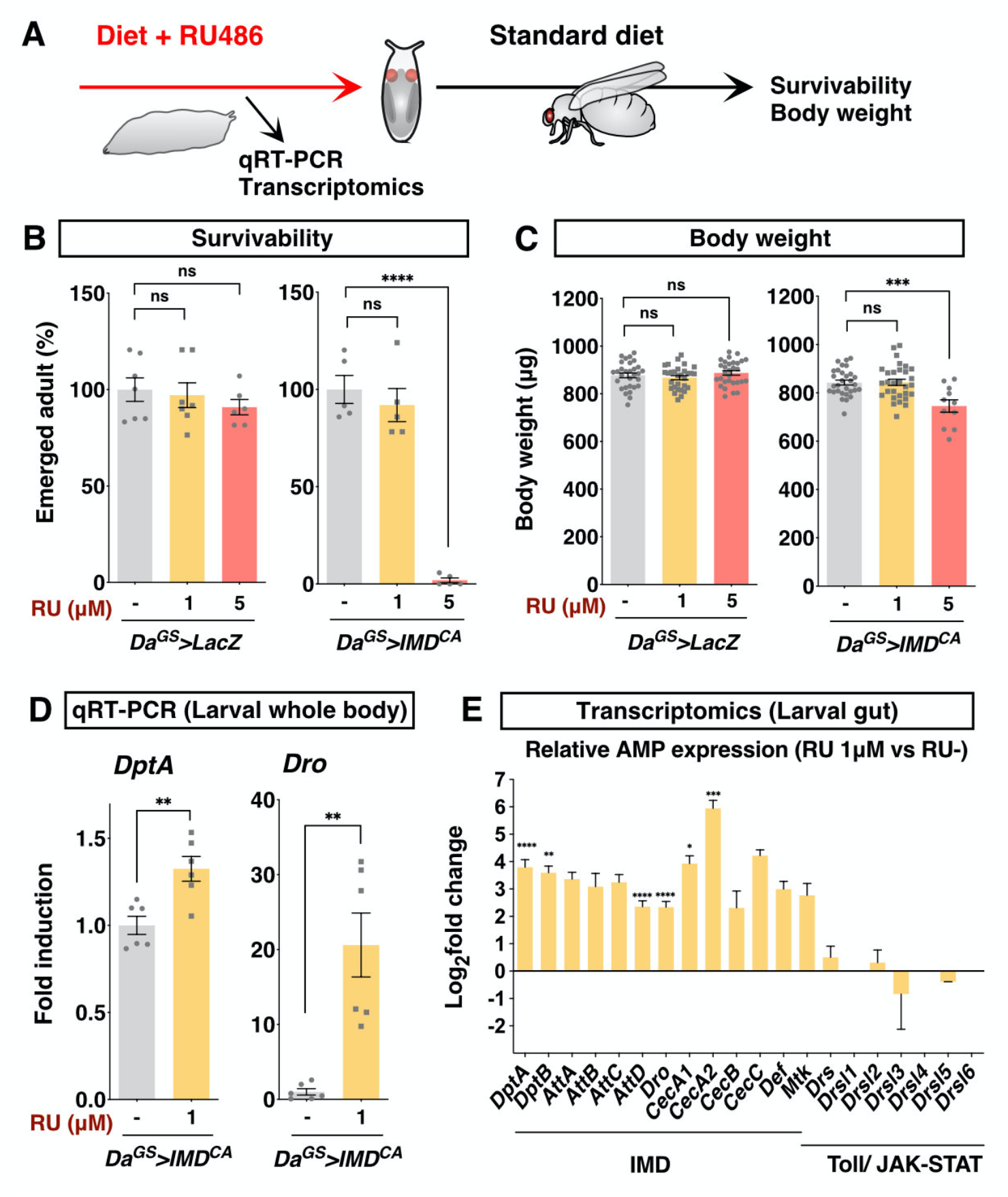
Mild IMD activation using Gene Switch (GS) system does not disturb development. (**A**), Experimental scheme. (**B**), Developmental survivability of flies expressing constitutive active form of IMD (*IMD^CA^*) or negative control (*LacZ*). Average numbers of flies successfully developed under each condition were shown. *Daughterless* Gene Switch driver is used to induce gene expression ubiquitously by RU486. n=5-7. (**C**), Adult body weight. One-week-old adult male flies are used. n=20-50 individual flies. (**D**), Quantitative RT-PCR of IMD target genes *Diptericin A* (*DptA)* and *Drosocin* (*Dro*) in the whole body of third instar larvae. n=6. (**E**), Transcriptomic analysis of the larval gut from *Da^GS^>IMD^CA^* third instar larvae. Relative to the negative control (no RU486 treatment) is shown. n=3. Each graph is shown as mean ± SEM. Statistics, **, *p*<0.01, ***, *p*<0.001, ****, *p*<0.0001, ns, not significant.

We first confirmed that gene expression was indeed induced with as low as 1 μM of RU486, as visualised by GFP expression (Fig. S1B). The driver activity is detected in the larval brain, the fat body, the gut and the Malpighian tubules (Fig. S1C). To quantify the level of IMD activation, we performed quantitative RT-PCR analysis for antimicrobial peptide (AMP) genes regulated by the IMD signalling pathway. IMD target genes *Diptericin A* (*DptA*) and *Drosocin* (*Dro*) were upregulated in the whole body of *Da^GS^>IMD^CA^* third instar larvae (Fig. 1D). These genes were upregulated mildly in various tissues such as the brain, the gut, and the fat body (Fig. S2A). We further performed a transcriptomic profiling by 3’mRNA-sequencing analysis using the gut tissue. AMPs predominantly regulated by the IMD pathway were all upregulated, while those regulated by other immune pathways were not, suggesting that IMD signalling was specifically activated in this experimental setting (Fig. 1E, Table S1). The list of differentially-expressed genes did not contain typical damage-responsive genes such as *upd3*, which is known to be massively increased in the larval gut upon oral infection of Ecc15 [19]. The AMP induction was already suppressed in the whole body of young adult male flies (Fig. S2B). Therefore, in this experimental setting, we can increase the IMD signalling pathway mildly in a juvenile-restricted manner.

### Larval immune activation influences adult fitness

The magnitude of IMD activation in the larvae of *Da^GS^>IMD^CA^* with 1 μM RU486 was weak enough to avert disturbance of developmental growth. We questioned whether this sublethal, transient immune activation in developing animals has a prolonged effect on adult physiology and ultimately alters lifespan (Fig. 2A). The lifespan of adult flies of *Da^GS^>IMD^CA^* fed with 1 μM of RU486 during the larval stage was significantly shortened in both male and female flies (Fig. 2B, C). In the control (*Da^GS^>LacZ*), RU486 did not affect the male lifespan at all, whereas it slightly decreased female lifespan. This side effect of RU486 on female flies would explain the greater extent in shortening female lifespan of *Da^GS^>IMD^CA^* with 1 μM of RU486 (Fig. 2C). We therefore decided to use male flies mainly for the rest of the study. The lifespan shortening by larval IMD activation was dose-dependent, as 2 μM RU486 further decreased the lifespan (Fig. 2B, C).

**Fig. 2.**
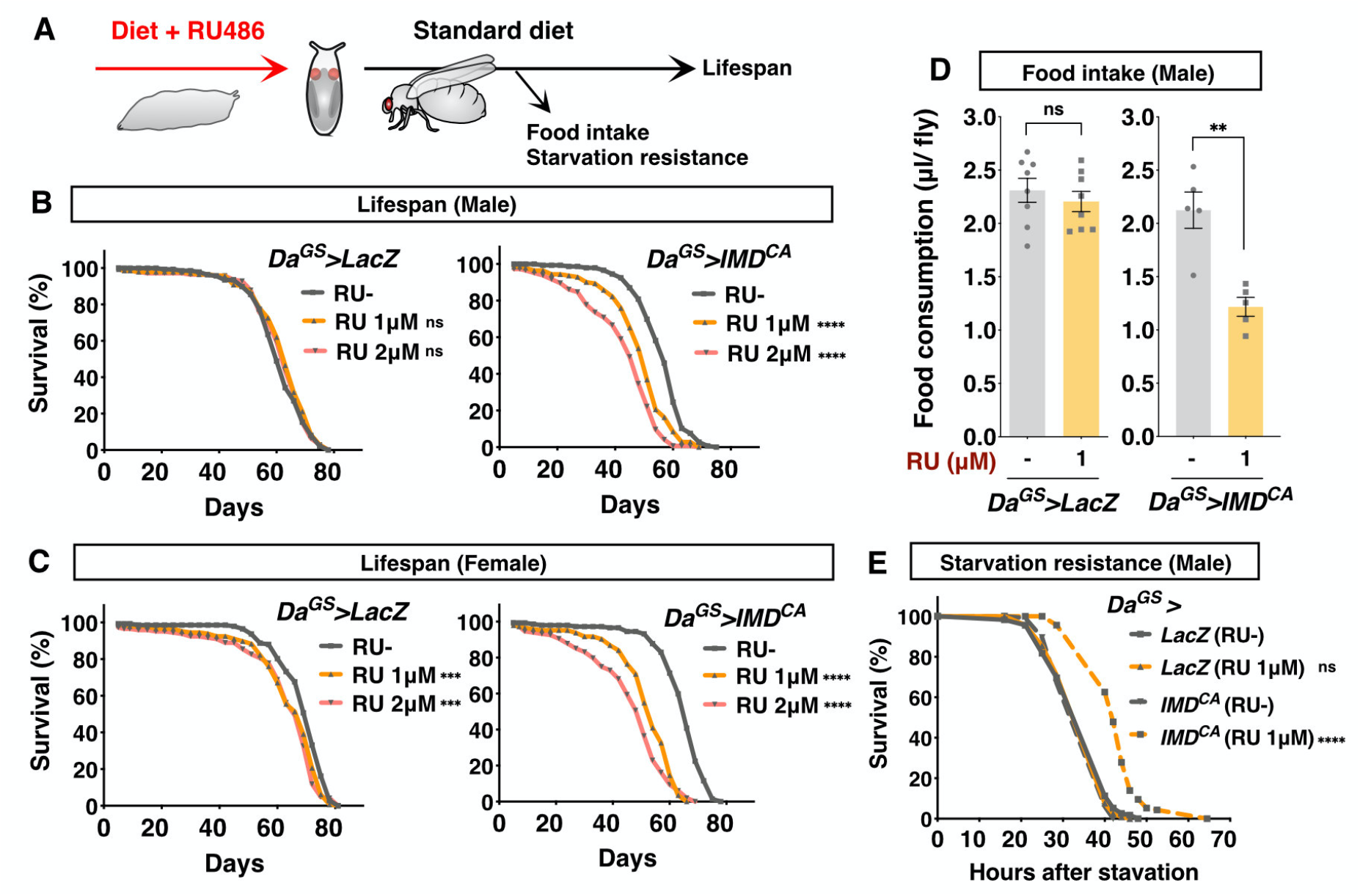
Larval IMD activation leads to shortened lifespan, decreased food intake, and increased starvation resistance in adult. (**A**), Experimental scheme. Food intake and starvation resistance were measured at one week after eclosion. (**B, C**), Lifespan of male (B) and female (C) flies with RU486 treatment during development. *Daughterless* Gene Switch driver (*Da^GS^*) was used to induce constitutive active form of IMD (*IMD^CA^*) or negative control (*LacZ*) ubiquitously by RU486 in the larval stage. n=120-150. (**D**), Amount of food intake of adult male flies assessed by capillary feeder (CAFE) assay. n=5-8 vials. (**E**), Survival curve under starvation condition (1% agar) of male flies t. n=70-80. Each graph is shown as mean ± SEM. Statistics, **, *p*<0.01, ***, *p*<0.001, ****, *p*<0.0001, ns, not significant.

To understand how adult fitness is altered by larval IMD activation, we further assessed the physiological conditions of the adult flies. Dietary protein amount is negatively correlated with lifespan. It was possible that the decrease of lifespan was owing to the increased food intake. Unexpectedly, young male flies that had experienced larval IMD activation took less food compared with the negative control (Fig. 2D). These data at least suggested that the lifespan shortening by larval IMD activation may not have been attributed to the dietary protein intake. Surprisingly, despite the decrease in food intake, they had increased starvation resistance (Fig. 2E). Larval IMD activation did not alter paraquat (oxidant) resistance, and it did induce hyper susceptibility to high salt stress (Fig. S3). These phenotypes invalidated the possibility that the shortened lifespan by larval IMD activation was simply due to the increased susceptibility to stresses (general sickness of the flies). Taken together, we concluded that IMD activation during development induces prolonged physiological changes in adult flies and decreases lifespan.

### Starvation resistance and lifespan are distinctively regulated by larval IMD activation

To identify which tissue(s) shortens lifespan upon IMD activation, we overexpressed *IMD^CA^* in a tissue-specific manner. We used Gene Switch drivers for neurons (*Elav^GS^*), the gut and the fat body (*TI^GS^*), and the Malpighian tubules (*Uro^GS^*) (Fig. 3A, Fig. S4). We observed that overexpression of *IMD^CA^* only by TI^GS^ decreased lifespan, suggesting that IMD activation in the larval gut and/or fat body induces shortened lifespan (Fig. 3B). Interestingly, however, starvation resistance was not elevated in *TI^GS^>IMD^CA^* flies, but this phenotype was rather observed in *Elav^GS^>IMD^CA^* flies (Fig. 3C). Therefore, shortened lifespan and starvation resistance are distinctive phenotypes triggered by the IMD activity from the different tissues. Similarly, the decreased food intake was induced only when *Elav^GS^* was used to drive IMD activation (Fig. 3D). The data indicate that food intake and starvation resistance have a correlation, while the lifespan shortening occurred in parallel. In this study, we focused on the lifespan phenotype. As the phenotype of lifespan shortening by *TI^GS^>IMD^CA^* is often weaker than *Da^GS^>IMD^CA^*, we used the *Da^GS^>IMD^CA^* for the rest of the study.

**Fig. 3.**
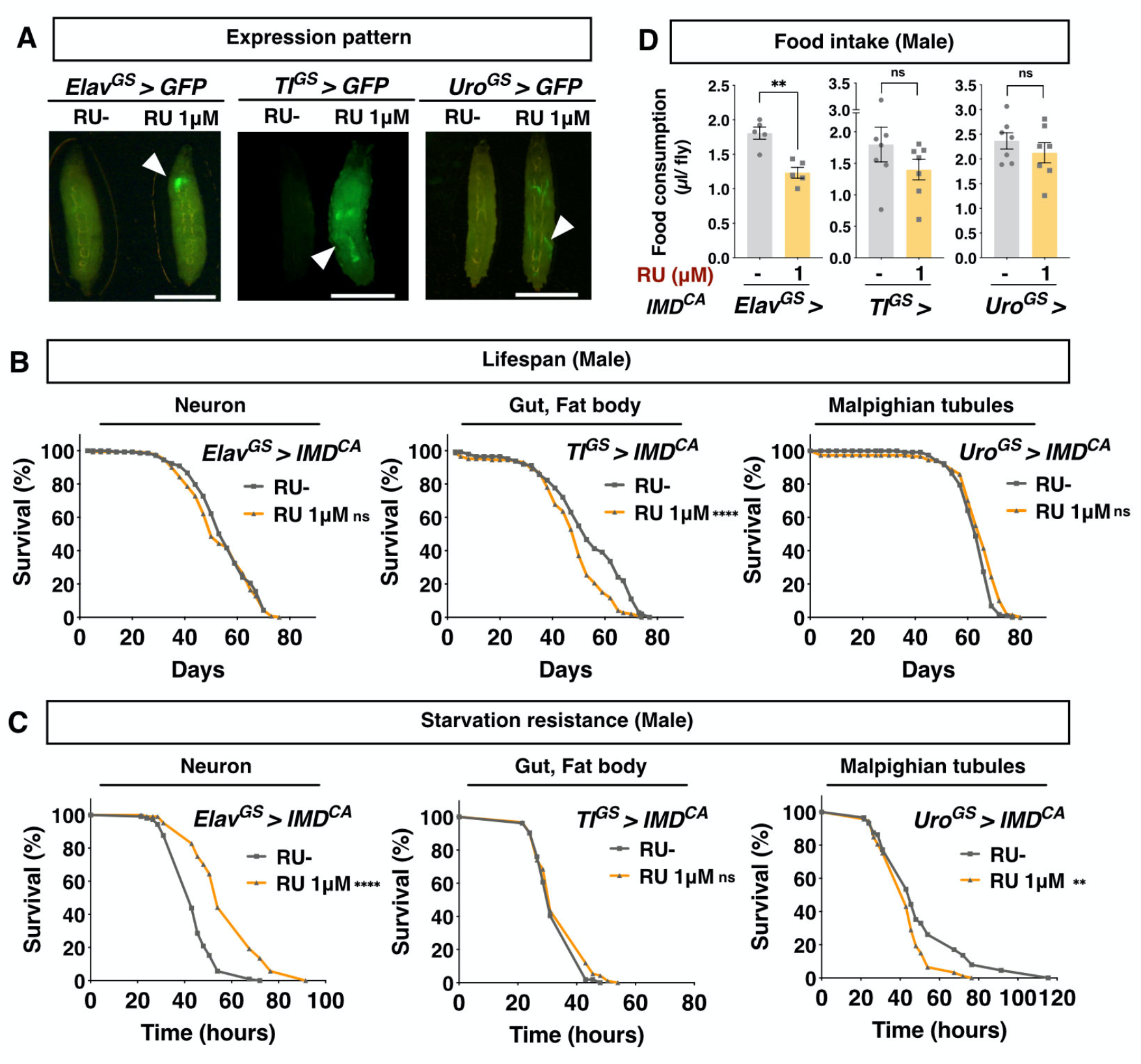
Tissue specific effect of larval IMD activation on adult phenotypes. (**A**), Expression pattern of each Gene Switch driver upon 1 μM of RU486. Arrowheads indicate the GFP expression. Scale bars, 1mm. (**B**), Lifespan of male flies. Gene Switch drivers are used to induce constitutive active form of IMD (*IMD^CA^*) by RU486 in the larval stage. n=120-150. (**C**), Survival curve of male flies under starvation condition (1% agar). n=70-80. (**D**), Amount of food intake of adult male flies assessed by capillary feeder (CAFE) assay. n=5-7 vials. Food intake and starvation resistance are measured at one week after eclosion. Each graph is shown as mean ± SEM. Statistics, **, *p*<0.01, ****, *p*<0.0001, ns, not significant.

### Larval IMD activation causes spontaneous immune activation in the adult gut

IMD activity is known to increase during ageing and negatively impacts organismal lifespan. To discover whether the shortened lifespan is due to the accelerated inflammatory response, we quantified the expression of *DptA* in the aged flies. We found that adult male flies with larval IMD activation showed an elevation of whole body IMD activity at five weeks of age (Fig. 4A). This IMD activity was likely derived from the gut as we detected a sharp increase of *DptA* expression in the gut (Fig. 4B).

**Fig. 4.**
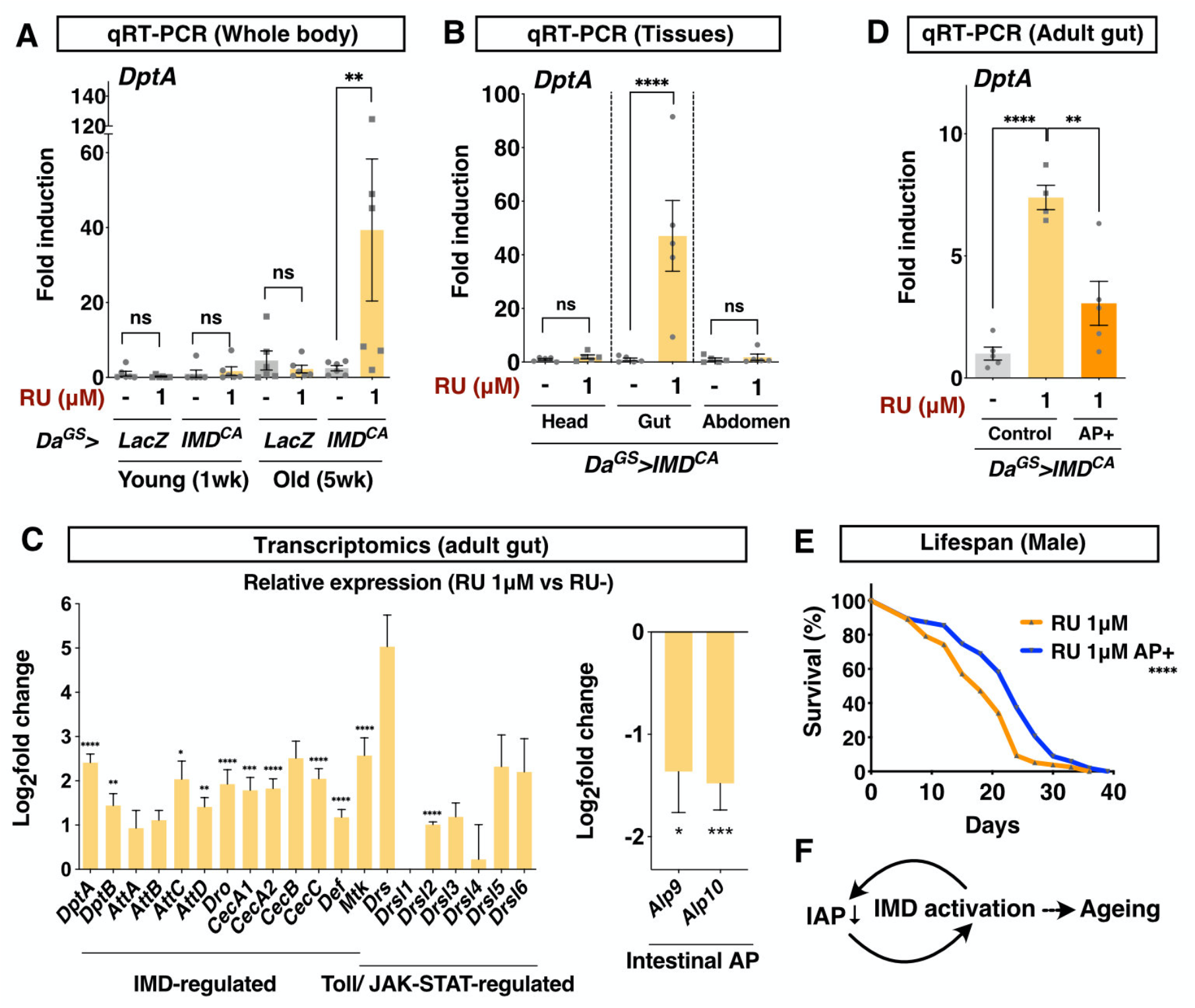
Larval IMD activation induces inflammatory intestine in adult. (**A**), Quantitative RT-PCR of IMD target gene *Diptericin A* (*DptA*) in the whole body of male flies. *Daughterless* Gene Switch driver (*Da^GS^*) is used to induce constitutive active form of IMD (*IMD^CA^*) or negative control (*LacZ*) ubiquitously by RU486 in the larval stage. n=6. (**B**), Quantitative RT-PCR of IMD target gene *Diptericin A* (*DptA*) in each body part of 5-week-old male flies. n=5. (**C**), Transcriptomic analysis of the adult gut from one-week-old *Da^GS^>IMD^CA^* male flies. Relative to the negative control (no RU486 treatment) is shown. n=3. (**D**), Quantitative RT-PCR of IMD target gene *Diptericin A* (*DptA*) in the adult gut. n=4-5. AP+, the flies fed with exogenous alkaline phosphatase supplemented on top of the fly food (100 unit/vial) for three days. (**E**), Lifespan of *Da^GS^>IMD^CA^* male flies with or without alkaline phosphatase supplementation (5 unit/vial). n=120-150. (**E**), Model. Each graph is shown as mean ± SEM. Statistics, *, *p*<0.05, **, *p*<0.01, ***, *p*<0.001, ****, *p*<0.0001, ns, not significant.

The fact that developmental IMD activation increased *DptA* expression in the adult gut suggested that an irreversible change occurred in this tissue. To describe the tissue condition, we performed a transcriptomic analysis of the young adult gut from *Da^GS^>IMD^CA^* fed with RU486 during the larval stage. Unexpectedly, on day 10 in adult male flies, many antimicrobial peptides were already upregulated (Fig. 4C, Table S2). Unlike the larval gut, where IMD target genes were specifically upregulated, the adult gut showed increased antimicrobial peptides regulated by Toll-or JAK/STAT as well (Fig. 4C). This observation suggested that increased antimicrobial peptides were not due to the sustained RU486 in their gut, but rather to the general inflammatory response of the tissue. Although we could not deny the possibility that overexpressed IMD^CA^ protein in the larval gut remained in the adult gut, it was less likely considering that the larval gut is completely degenerated and replaced by the newly-generated adult gut [23-25].

We assumed that an experience of larval immune activation augments immunity as an adaptive response to prepare for future infection in the adult flies. As the increased *DptA* expression is restricted in the gut, we asked whether the flies were resistant to oral infection that could be influenced by AMPs [26]. Unexpectedly, the larval IMD induction was not beneficial for adult flies against *Pseudomonas entomophila* infection, but rather it increased susceptibility (Fig. S5). Therefore, higher IMD activity in the gut is thought to be pathological and implies accelerated tissue ageing.

We also noticed that intestinal alkaline phosphatase (IAP) *Alp9* and *Alp10* were decreased in the adult gut (Fig. 4C, Table S2). Among 18 downregulated genes (Fold change<0.5, *p*<0.05), two IAPs were listed in the third and fourth place, the expression of which decreased by one third of the control. IAP is an evolutionally-conserved regulator of gut homeostasis, the expression of which is known to be downregulated during ageing [27]. Decreased IAP expression is also reported in rodents and in human patients of inflammatory bowel disease [28]. Increased IAP expression is beneficial to suppress DSS-induced colitis in mice [29]. Exogenous Alp supplementation can augment the tissue homeostasis and extend mouse and fly lifespans [27]. In our model, feeding adult flies that experienced larval IMD activation with a high dose of Alp (100 units/vial) suppressed *DptA* upregulation (Fig. 4D). Sustained Alp (5 units/vial) supplementation increased the lifespan (Fig. 4E). These data suggest that decreased IAP expression, at least partially, contributes to the inflammatory response in the gut and the concomitant shortened lifespan. Decreased IAP expression did not occur in the larval gut (Table S1). Alp10 was intriguingly listed in the upregulated genes in the larval gut. Genetic IMD activation in the adult gut decreased IAP expression, suggesting that decreased IAP is likely due to the upregulated IMD signalling (and/or concomitant tissue senescence) (Fig. S6). Considering that IAP suppresses IMD activity, developmental IMD activation could trigger the inflammatory vicious cycles of IAP downregulation and IMD upregulation (Fig. 4F).

### Gut microbiota exacerbate gut immune activation and shortened lifespan

Gut microbiome increases the innate immune activation during ageing, which shortens lifespan [4,30]. *Acetobacteraceae*, such as *Acetobacter aceti*, is known to increase IMD activity in aged flies, and removing it results in extended lifespan [5]. Another *Acetobacteraceae Gluconobacter* spp. are known to expand in the gut microbiota in response to the host immune activation, and induce mortality of the flies [31,32]. We assumed that altered gut microbiota might contribute to the lifespan shortening in adult flies that experienced larval IMD activation. To test this hypothesis, we performed 16S rRNA gene amplicon sequencing analysis of the gut microbiota in the young adult gut. The result did not delineate a huge difference in the microbial composition (Fig. 5A). The total number of live bacteria assessed by colony forming unit assay was also not significantly changed (Fig. 5B). However, when we quantified the number of bacteria by quantitative PCR using a set of primer detecting genera *Acetobacter* or *Gluconobacter*, we noticed that *Gluconobacter* was significantly increased (Fig. 5C). Although we cannot distinguish whether this dysbiosis would be a consequence or a cause of the immune response, the change of the gut microbiome is another evidence that the flies suffer from the inflammatory intestine.

**Fig. 5.**
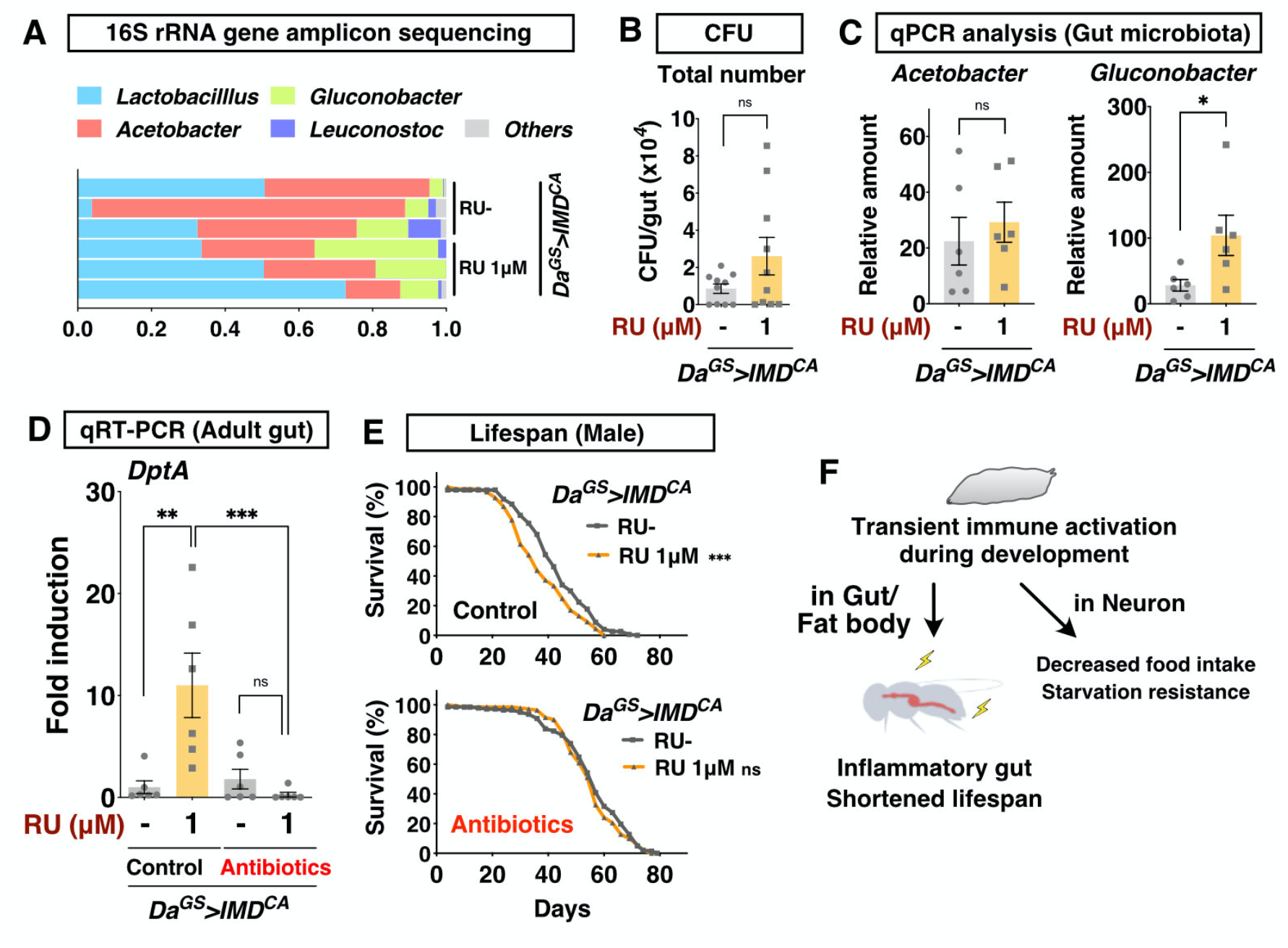
Removing gut microbiota rescues shortened lifespan. (**A**), 16S rRNA gene amplicon sequencing analysis of adult male gut. Each genus is shown using different color. Three biological replicates for each condition. *Daughterless* Gene Switch driver (*Da^GS^*) is used to induce constitutive active form of IMD (*IMDCA*) ubiquitously by RU486 in the larval stage. (**B**), Colony forming unit assay to count the number of live bacteria from the adult male gut. n=10. (**C**), Quantitative PCR of *Acetobacter* or *Gluconobacter* in the adult male gut. n=6. (**D**), Quantitative RT-PCR of IMD target gene *Diptericin A* (*DptA*) in the adult gut. See methods for the composition of antibiotic cocktail. n=6. These phenotypes are analysed at one week after eclosion (**A-D**). (**E**), Lifespan of male flies with or without antibiotics supplementation in adult. n=120-150. (**F**), Model. Each graph is shown as mean ± SEM. Statistics, *, *p*<0.05, **, *p*<0.01, ***, *p*<0.001, ns, not significant.

To discover whether the gut microbiota are involved in the gut inflammatory response and eventually shorten lifespan, we fed adult flies that experienced larval IMD activation with antibiotics (Rifampicin, Tetracycline, and Ampicillin) to eliminate gut microbiota. We found that IMD upregulation in the adult gut was abolished by the antibiotic treatment (Fig. 5D). In this condition, larval IMD activation did not shorten lifespan (Fig. 5E). These data suggest that gut microbiota contributed to the pathological phenotypes. Together, larval IMD activation promotes the dysregulated gut homeostasis including both host immune response and the gut microbiota (Fig. 5F).

## Discussion

In this study, we developed a model to study how developmental immune activation influences adult fitness and ultimately organismal lifespan. Strong activation of the immune signalling pathway inhibits the developmental processes and causes lethality [33,34]. By utilising the Gene Switch system, we set up juvenile-restricted low-grade IMD activation that enabled us to obtain superficially-normal, healthy adult flies. Nonetheless, this larval IMD activation led to the development of the adult gut with induced antimicrobial peptide genes, decreased *Alp* expression, and increased *Gluconobacter* sp. in the gut microbiota, all of which are hallmarks of inflammatory gut. We assume that transient IMD activation in the larval gut sensitises the adult gut to inflammatory cues. This phenomenon might be analogous to “trained immunity” or innate immune memory by which greater protection against reinfection is achieved [35,36]. We could not observe, however, any protection of adult flies that experienced larval IMD activation to oral infection despite them having increased antimicrobial peptides. This might suggest that the chronic immune activation in the adult gut is pathological and damages the gut epithelium rather than protecting it.

Mechanistically, it is possible that the sensitivity of IMD signalling becomes high partly due to the dampened expression of a negative regulator of IMD, such as IAPs. We previously observed that the systemic inflammatory response in necrosis-induced flies decreases S-adenosylmethionine (SAM), a methyl donor required for histone methylation [37]. In worms, SAM is decreased during infection, and this leads to decreased H3K4me3 to regulate immune response [38]. Spontaneous immune response in aged fat body in *Drosophila* is attributed to declined lamin C expression and epigenetic deregulation [39]. Therefore, the immune-epigenetic crosstalk in the adult midgut progenitors in the larval gut would consequently alter the epigenetic homeostasis of the adult intestinal stem cells and/or their progeny enterocytes. The detailed mechanism of the IMD sensitisation needs to be further investigated.

It is widely accepted that early-life exposures to microorganisms, such as healthy gut microbiota, are essential for shaping appropriate immune and metabolic homeostasis [40-43]. Inappropriate microbial exposures therefore impact various inflammation-associated diseases in later life, including inflammatory bowel disease [44,45]. Mice born of germ-free mothers become susceptible to a high-fat diet inducing obesity, due to the loss of immune, endocrinal homeostasis developed in the absence of bacterial metabolites [46]. Early-life disturbance of microbial composition by transient antibiotic treatment can cause obesity in adults [47,48]. Antibiotic treatment during development can also induce long-term changes in cytokine production in the brain and associated behavioural alteration [49]. Whether immune signalling *per se* in the developmental stage provokes inflammatory diseases and affects organismal lifespan, as we observed in flies, needs to be tested in mammals.

The important question raised by the present study is how neuronal immune activation in the larval stage leads to starvation resistance and decreased food intake in adults. Infection by pathogen decreases food intake, at least in larvae [26]. Direct IMD activation by circulating peptidoglycan in octopaminergic neurons alters oviposition but does not affect food intake [50]. As far as we know, it has not been elucidated whether activation of IMD signalling in some neurons regulates food intake in adult flies. Increased starvation resistance suggests decrease in energy expenditure and/or augmented metabolic efficiency of the animals. Immune-metabolic switch is essential for the allocation of nutrients from anabolism to immune effector production [51]. Persistent activation of the IMD pathway in the fat body leads to altered metabolism [52]. IMD activation by gut microbiota can also modulate metabolic homeostasis of the host [53,54]. IMD activation in enteroendocrine cells in the gut alters the metabolism and development through an endocrine peptide Tachykinin [55]. In our model, we assume that experiencing IMD activation triggers the thrifty phenotype by modulating metabolism in the animals to prepare for future infection/stress response. This long-term immuno-metabolic interaction is an interesting direction to be explored in a future mechanistic study.

## Methods

### Drosophila stocks and husbandry

Flies were reared on a standard diet containing 4% cornmeal, 6% baker’s yeast (Saf Yeast), 6% glucose (Wako, 042-31177), and 0.8% agar (Kishida Chemical, 260-01705) with 0.3% propionic acid (Tokyo Chemical Industry, P0500) and 0.05% nipagin (Wako, 132-02635). Flies were reared under 25°C, 65% humidity with 12 h/12 h light/dark cycles. The fly lines were: *Da-GeneSwitch* [56], *UAS-lacZ* (Gifted from Dr. Corey S. Goodman), *UAS-IMD^CA^* [6], *Elav-GeneSwitch* [22], *TI-GeneSwitch* (Gifted from Dr. Scott Pletcher), *Uro-GeneSwitch* (This study), *UAS-2×EGFP* (Bloomington 6874). *Da-GeneSwitch, UAS-lacZ* and *UAS-IMD^CA^* were backcrossed eight generations with *w^iso31^*. Embryos were collected using a cage containing young (roughly 1-week-old) parents of GS and UAS lines and an acetic acid agar plate (2.3% Agar (BD214010), 10% Sucrose (Wako 196-00015), 0.35% Acetic acid (Wako 017-0256)) with live yeast paste. Equal volumes of the collected embryos were put onto the top of fly food containing RU486 (Tokyo Chemical Industry, M1732, dissolved in ethanol) or its negative control ethanol, in order to control the larval density. Adult flies eclosed within two days were collected and maintained for additional two days for maturation on standard fly diet. Then, flies were sorted by sex, put 15 flies per vial, and flipped to fresh vials every three days. Antibiotics (200 μg/mL rifampicin, 50 μg/mL tetracycline, 500 μg/mL ampicillin, together with 0.12% nipagin) were added to standard diet to remove all bacteria.

### Developmental speed, survivability, and body weight measurement

After putting the equal amount of embryos, the number of pupae on each time point was counted for developmental speed. Data were normalised by the total number of pupae. For survivability, the number of adult flies for each bottle was counted, and divided by the number of the control (without RU486 treatment). The body weight of eclosed adult flies was measured individually by an ultra-microbalance (METTLER TOLEDO, XPR2).

### Construction of *Uro-GeneSwitch* fly

To generate the *Uro-GeneSwitch* driver, the putative 881 bp promotor sequence of *Urate oxidase* (*Uro*) gene was amplified by PCR using *w^Dah^* genomic DNA. The sequence of GeneSwitch was amplified by PCR using pElav-GeneSwitch (Addgene: 83957). The backbone vector pElav-GeneSwitch was digested with KpnI, and ligated with the amplicons Uro promoter and GeneSwitch using NEBuilder HiFi DNA Assembly Kit (NEW ENGLAND BioLabs, E2621X). Transgenic lines were generated using standard methods for P-element-mediated germline transformation (BestGene Inc).

### RNA sequencing analysis

Dissected larval or adult guts were homogenised in 150 μL of QIAzol Lysis Reagent (QIAGEN, 79306), and stored at −80 °C. Triplicate was prepared for each experimental group, containing three to five male guts per sample. Then, 350 μL of QIAzol Lysis Reagent was added and left for 30 minutes at room temperature. 100 μL of chloroform was added and mixed with vortex, then left for two minutes at room temperature. Using RNeasy Plus Micro Lit (QIAGEN, 74034), RNA was extracted based on manufacturer’s protocol. RNA was sent to Kazusa Genome Technologies to perform 3’ RNA-seq analysis. cDNA library was prepared using QuantSeq 3’ mRNA-Seq Library Prep Kit for Illumina (FWD) (LEXOGEN, 015.384). Sequencing was done using Illumina NextSeq 500 and NextSeq 500/550 High Output Kit v2.5 (75 Cycles) (Illumina, 20024906). Raw reads were analysed by BlueBee Platform (LEXOGEN), which performs trimming, alignment to *Drosophila* genome, and counting of the reads. The count data was statistically analysed by Wald test using DESeq2. The result has been deposited in DDBJ under the accession number DRA011490.

### Quantitative RT-PCR analysis

Total RNA was purified from five male flies or three-to-five guts using Promega ReliaPrep RNA Tissue Miniprep kit (z6112). cDNA was made from 200 or 400 ng DNase-treated total RNA by the Takara PrimeScript RT Reagent Kit with gDNA Eraser (RR047B). Quantitative PCR was performed using TB Green™ Premix Ex Taq™ (Tli RNaseH Plus) (Takara bio RR820W) and a Quantstudio6 Flex Real Time PCR system (ThermoFisher) using *RNA pol2* as an internal control. Primer sequences were listed in Table S3.

### 16S rRNA gene amplicon sequencing analysis and quantitative PCR of bacteria

Adults were briefly rinsed in PBST (0.1% Triton X-100), 50% (v/v) bleach (OYALOX), 70% ethanol, and PBS before dissection. Male guts (6-8 per sample) without tracheae, Malpighian tubules and crop were dissected from day 10 adult flies. Dissected guts were collected in PBS on ice, then homogenised in 270 μL of lysis buffer (20 mM Tris pH8.0, 2 mM EDTA and 1% Triton X-100) with 20 mg/mL lysozyme from chicken egg (Sigma, L4919) using a tissue grinder (BMS BC-G10) with a pestle (BMS BC-PES50S). The homogenates were incubated at 37°C for 45 min in a 1.5 mL microcentrifuge tube, then further homogenised in a 2 mL tube (Yasui Kikai, ST-0250F-O) containing 0.1 mm glass beads (Yasui Kikai YZB01) using a Multi-beads shocker (Yasui Kikai) at 2500 rpm for 20 sec ×2. To remove bubbles, the tube was centrifuged briefly. After an additional 15 min incubation at 37°C, 30 μL of proteinase K and 200 μL of Buffer TL (Qiagen) were added to each sample. The samples were incubated at 56°C for 15 min. Genomic DNA was purified by QIAamp DNA Micro kit (Qiagen, 56304) and sent to Macrogen Corp. Japan where 16S rRNA amplicon sequencing (Illumina MiSeq) and the bioinformatics analysis was performed. 16S rRNAs were amplified using primers targeting V3 and V4 region. The result has been deposited in DDBJ under the accession number DRA011489.

For quantification of bacterial species by quantitative PCR, three different primer sets were used for *Acetobacter* [57], *Gluconobacter* [58] and *Drosophila GAPDH* gene for normalisation. Primer sequences were listed in Table S3. For *Acetobacter*, TB Green™ Premix Ex Taq™ (Tli RNaseH Plus) (Takara bio RR820W) was used. For the analysis of *Gluconobacter*, probe-based quantitative PCR was performed using PrimeTime Gene Expression Master Mix (Integrated DNA Technologies, 1055772).

### Colony Forming Unit assay

One fly from each vial is dissected in PBS and homogenised in PBS. The serial dilutions of the homogenate were plated by EDDY JET2 (iUL, PL0300) and the number of colonies on MRS agar plate (Kanto Chemical, 711361-5) was counted after incubation for two days at 30°C.

### Capillary feeder assay for food intake

Two glass capillaries containing 5% sucrose, 2 mg/mL red dye (Acid red 52, Wako, 3520-42-1), and n-Octyl acetate (1:100,000; TCI, 112-14-1) were inserted into the cap. Ten male flies were placed in each vial containing 1% agar to avoid desiccation stress. The level of the food was marked, and the vials were laid in a container with wet towels to prevent water evaporation. The container was incubated at 25°C. After 24 hours, the amount of the food remained in the capillaries was recorded. The vial without flies was also included in the container to subtract evaporation.

### Survival assays

For lifespan analysis, the number of dead flies was counted every three days. To minimise the variation between culturing vials, we used 8-12 vials with 15 flies/vial. For high salt stress, flies were placed onto food containing 500 mM NaCl (Wako, 191-01665), 5% sucrose (Wako, 196-00015), 1% agar (Kishida Chemical, 260-01705). For starvation stress, flies were placed in vials containing 1% agar. For oxidative stress, flies were placed onto food containing 10 mM paraquat (1,1’-Dimethyl-4,4’-bipyridinium Dichloride, Tokyo Chemical Industry, D3685), 5% sucrose, 1% agar. In each assay 15 male flies/vial were incubated at 25°C and the number of dead flies was counted several times in a day.

*P. entomophila* wild-type strain L48 was kindly provided by Dr. B. Lemaitre. Oral infection was performed as described previously [59]. Briefly, *P. entomophila* was grown in LB medium at 29°C overnight and collected by centrifugation. Adult flies were incubated for 2 h at 29°C in an empty vial for starvation and then placed in a fly vial with a bacterial solution. The bacterial solution was obtained by mixing a pellet of bacteria with a culture supernatant (1:1), added to a filter disk that completely covered the surface of the standard fly medium. Flies were maintained at 29°C and mortality was monitored.

### Intestinal alkaline phosphatase supplementation

5,000 units/mL Quick CIP (New England Lab, M0525L) was diluted by enzyme buffer (25 mM Tris-HCl, 1 mM MgCl_2_, 0.1 mM ZnCl_2_, 50% Glycerol, pH=7.5, 25°C). Twenty microlitters of the CIP solution were directly applied on top of the standard diet. The enzyme buffer was used as the negative control. For quantification of *DptA* expression, the adult flies were fed with 100 units/vial for three days. For lifespan analysis, only 5 units/vial was used for economical reason to treat adult flies throughout the life.

### Quantification and statistical analysis

Statistical analysis was performed using Graphpad Prism 8 except for survival curves where OASIS2 was used [60]. A two-tailed Student’s *t*-test was used to test between two samples. One-way ANOVA with Sidak’s test was used to compare any combination of interest within a group. Statistical significance; *, *p*<0.05, **, *p*<0.01, ***, *p*<0.001, ****, *p*<0.0001. Bar graphs were drawn as mean and SEM with all the data point shown by dots to allow readers to see the number of samples and each raw data.

## Acknowledgments

We thank Corey Goodman, Edan Foley, Bruno Lemaitre, and Scott Pletcher for *Drosophila* stocks. We thank Yoriko Akuzawa-Tokita for technical assistance and all other members of Miura’s laboratory for active discussion. This work was supported by AMED-PRIME to F.O. under Grant Number 20gm6010010h0004 and 20gm6310011h0001 and to T.K. under Grant Number 20gm6010011h0004, and by AMED-Project for Elucidating and Controlling Mechanisms of Aging and Longevity to M.M under Grant Number JP20gm5010001. This work was also supported by grants from the Japan Society for the Promotion of Science to F.O. under Grant Number 19H03367 and 20H05726, and to M.M. under Grant Number 16H06385.

## Author contributions

F.O. and M.M. conceived and supervised the project. K.Y. and A.O. performed experiments and analysed data. H.Kosakamoto established methodology and analysed data. T.Y. established UroGS fly and assisted some experiments. H.Kadoguchi and T.K. performed oral infection experiment and suggested directions. F.O. wrote the initial manuscript. All authors edited and approved the final manuscript.

## Supplementary Figures and Tables

**Fig. S1.**
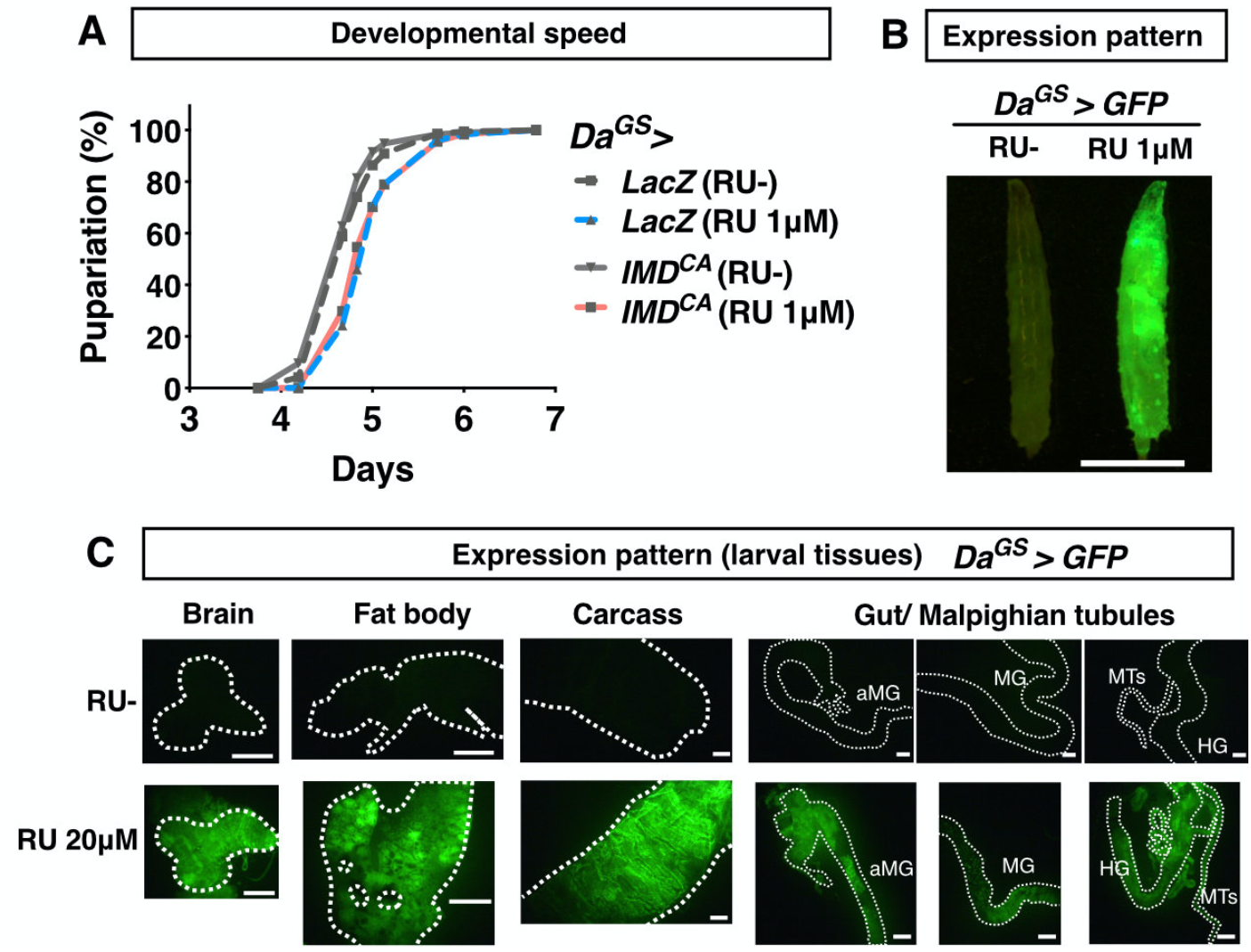
Mild IMD activation using Gene Switch system does not delay development. (**A**), Developmental timing of flies expressing constitutive active form of IMD (*IMD^CA^*) or negative control (*LacZ*) by *Daughterless* Gene Switch driver (*Da^GS^*) with or without 1 μM of RU486. (**B**), Expression pattern of *Daughterless* Gene Switch driver upon 1 μM of RU486 visualised by GFP. Scale bar, 1mm. (**C**), Expression pattern of *Daughterless* Gene Switch driver in the larval tissues. GFP is induced by 20 μM of RU486. Scale bars, 200 μm.

**Fig. S2.**
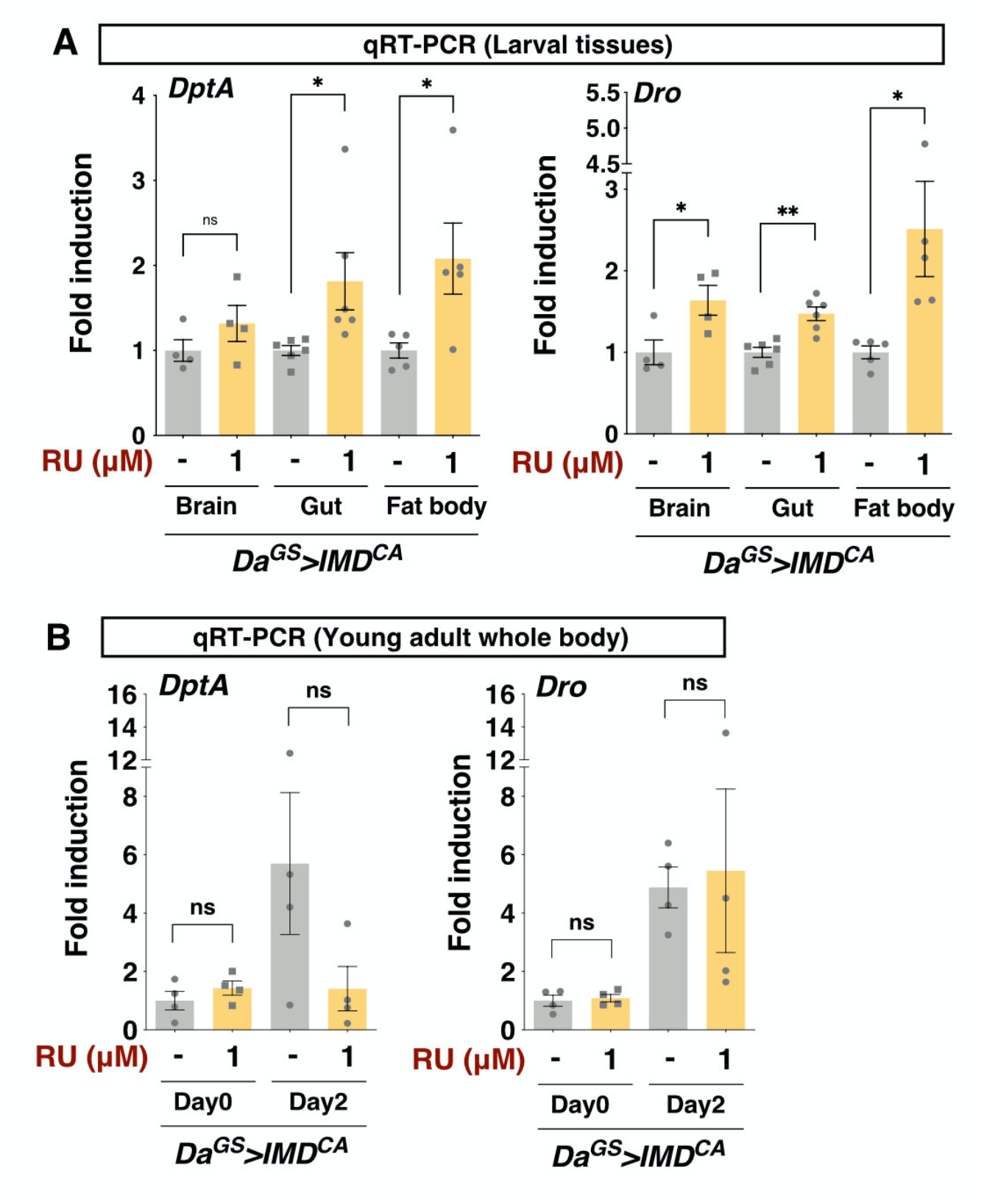
Transient Larval IMD activation by *Da^GS^*. (**A**), Quantitative RT-PCR of IMD target genes *Diptericin A* (*DptA*) and *Drosocin* (*Dro*) in the third instar larval organs. *Daughterless* Gene Switch driver (*Da^GS^*) is used to induce constitutive active form of IMD (*IMD^CA^*) or negative control (*LacZ*) ubiquitously by RU486. n=4-6. (**B**), Quantitative RT-PCR of IMD target genes *Diptericin A* (*DptA*) and *Drosocin* (*Dro*) in the whole body of day0 or day2 male flies. n=4. Each graph is shown as mean ± SEM. Statistics, *, *p*<0.05, **, *p*<0.01, ns, not significant.

**Fig. S3.**
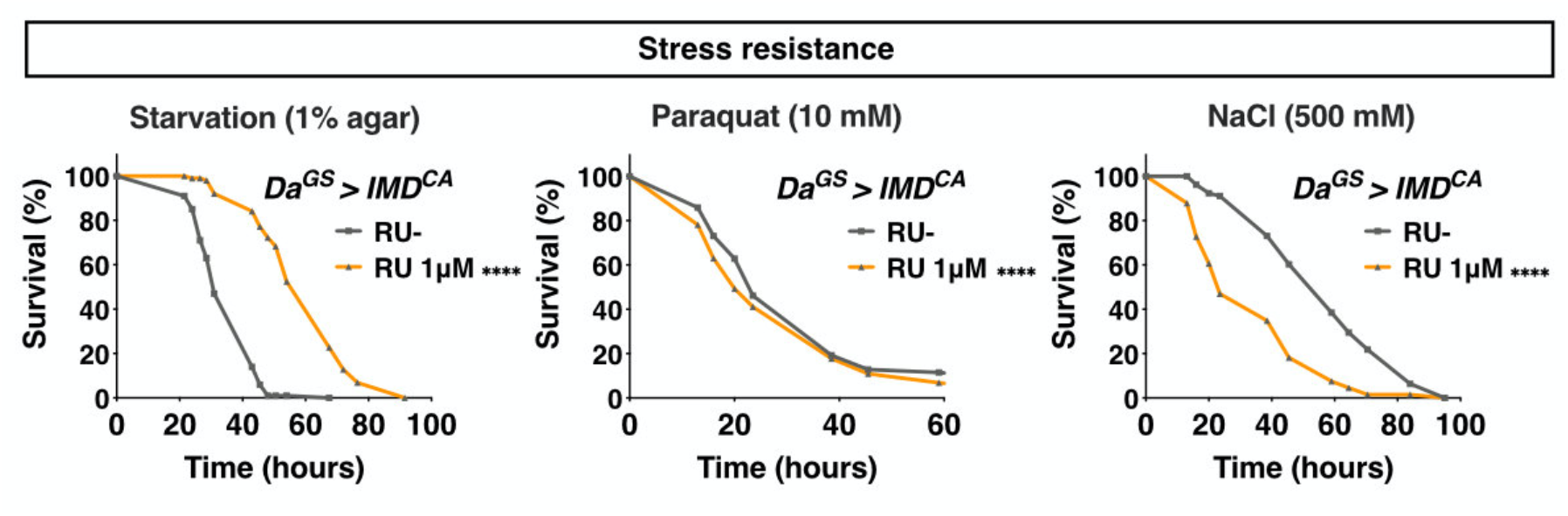
Larval IMD activation alters resistance to environmental stressors in adult. (**A**), Survival curves under starvation (1% agar), paraquat (10 mM), or high-salt (NaCl 500 mM) conditions of one-week-old male flies that experienced larval IMD activation. *Daughterless* Gene Switch driver (*Da^GS^*) is used to induce constitutive active form of IMD (*IMD^CA^*) ubiquitously by RU486 in the larval stage. n=70-80.

**Fig. S4.**
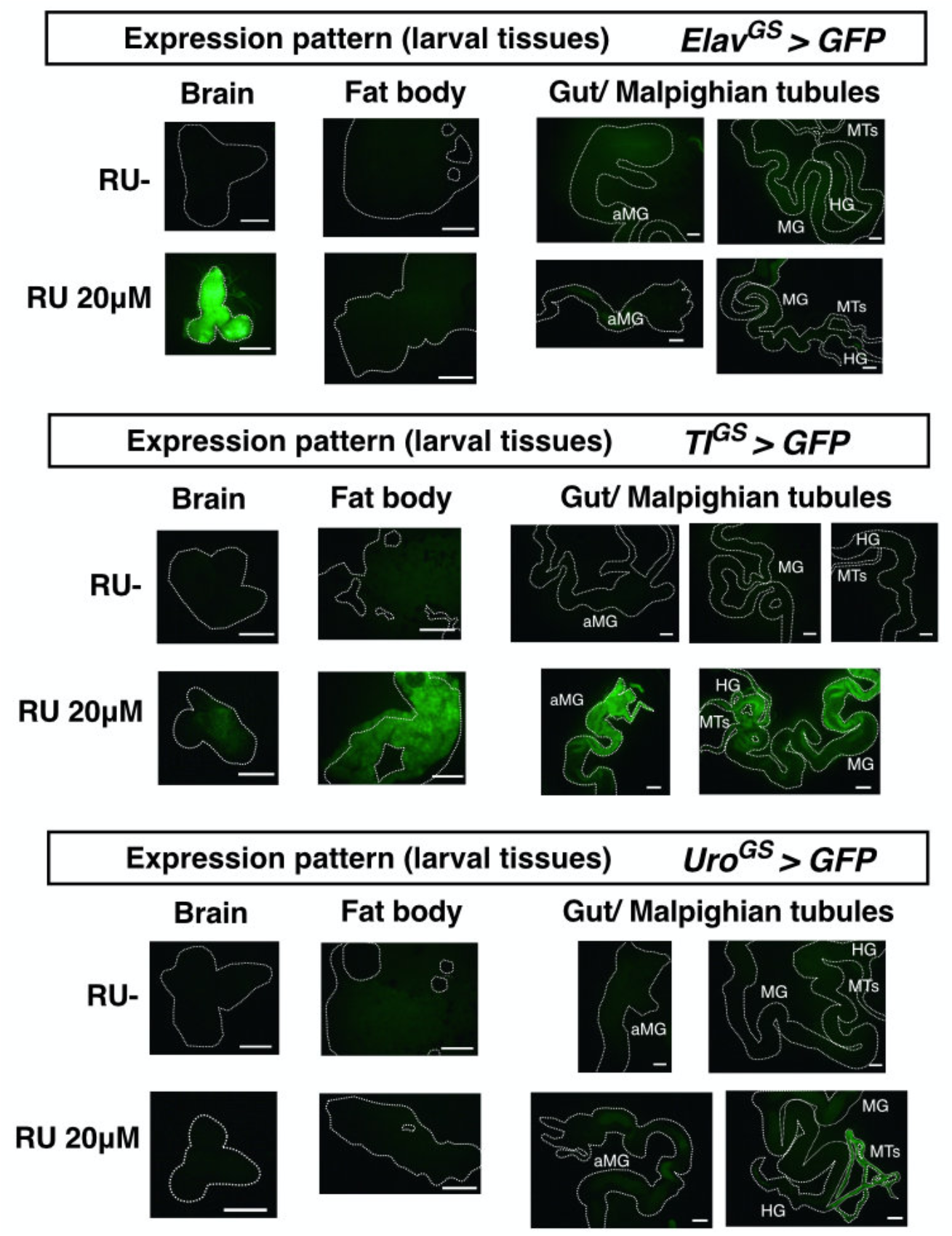
Expression patterns of Geneswitch drivers in the larval tissues. GFP was expressed by *Elav^GS^, TI^GS^*, and *Uro^GS^* drivers using 20 μM RU486. Third-instar larvae were dissected and GFP was visualised by fluorescent microscopy. Scale bars, 200 μm.

**Fig. S5.**
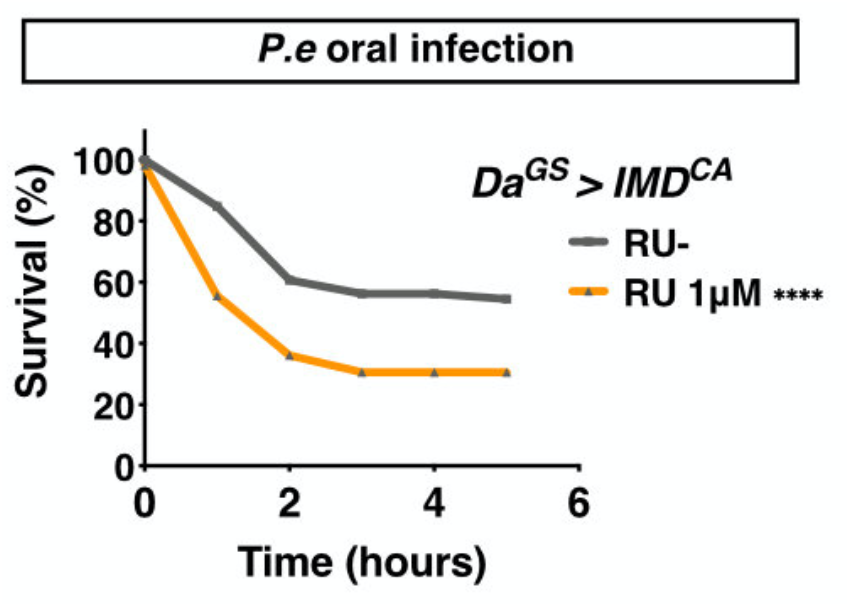
Adult flies upon larval IMD activation are susceptible to oral infection. Survival curves upon *P. entomophila* oral infection of one-week-old male flies that have experienced larval IMD activation. *Daughterless* Gene Switch driver (*Da^GS^*) is used to induce constitutive active form of IMD (*IMD^CA^*) ubiquitously by RU486 in the larval stage. n=100-110.

**Fig. S6.**
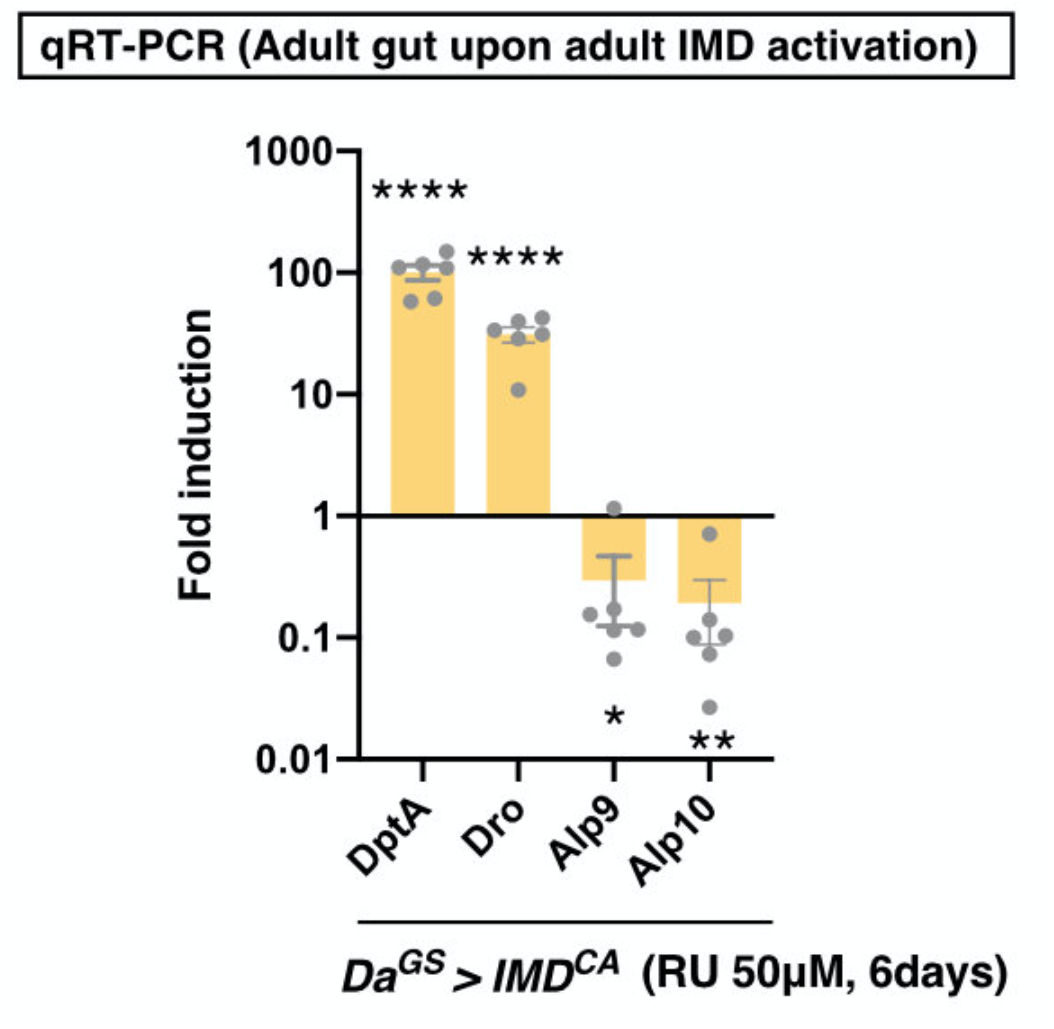
IMD activation in adult flies decreases IAP expression. Quantitative RT-PCR of *DptA, Dro, Alp9*, and *Alp10* in the adult gut. *Daughterless* Gene Switch driver (*Da^GS^*) is used to induce constitutive active form of IMD (*IMD^CA^*). *Da^GS^>IMD^CA^* female flies were fed with 50 μM RU486 for 6 days. n=6. Each graph is shown as mean ± SEM. Statistics, *, *p*<0.05, **, *p*<0.01, ****, *p*<0.0001, ns, not significant.

**Table S1.** Differentially-expressed genes in the larval gut upon mild IMD activation (Fold change>1.5 or <0.67, adjusted *P*-value<0.05).

| Symbol | Gene Name | Fold Change | *p*-value |
| --- | --- | --- | --- |
| **Up-regulated genes** |  |  |  |
| CecA2 | Cecropin A2 | 61.82 | 4.91.E-04 |
| CecA1 | Cecropin A1 | 15.21 | 2.78.E-02 |
| DptA | Diptericin A | 13.86 | 6.27.E-12 |
| DptB | Diptericin B | 12.12 | 1.41.E-03 |
| Zip42C.1 | Zinc/iron regulated transporter-related protein 42C.1 | 11.41 | 5.30.E-03 |
| AttD | Attacin-D | 5.12 | 2.46.E-06 |
| Dro | Drosocin | 5.05 | 1.69.E-05 |
| edin | elevated during infection | 4.76 | 1.68.E-02 |
| CG14606 | uncharacterized protein | 3.64 | 6.30.E-03 |
| CG14205 | uncharacterized protein | 3.02 | 1.97.E-03 |
| CG1139 | uncharacterized protein | 2.73 | 7.28.E-03 |
| l(2)34Fc | lethal (2) 34Fc | 2.50 | 6.03.E-03 |
| Diedel3 | Diedel 3 | 2.42 | 3.28.E-03 |
| CG13078 | uncharacterized protein | 2.38 | 4.84.E-02 |
| CG32751 | uncharacterized protein | 2.31 | 6.98.E-04 |
| CG13325 | uncharacterized protein | 2.29 | 1.51.E-04 |
| CG5157 | uncharacterized protein | 2.26 | 2.26.E-03 |
| CG15255 | uncharacterized protein | 2.15 | 1.51.E-02 |
| Amy-d | Amylase distal | 1.97 | 6.03.E-03 |
| Ugt37A2 | UDP-glycosyltransferase family 37 member A2 | 1.94 | 5.23.E-03 |
| CG17570 | uncharacterized protein | 1.91 | 5.23.E-03 |
| Pdxk | Pyridoxal kinase | 1.83 | 1.41.E-03 |
| CG8773 | uncharacterized protein | 1.76 | 1.23.E-02 |
| CG4752 | uncharacterized protein | 1.70 | 2.75.E-02 |
| JhI-26 | Juvenile hormone-inducible protein 26 | 1.68 | 2.75.E-02 |
| Npc2c | Niemann-Pick type C-2c | 1.65 | 2.41.E-02 |
| Alp10 | Alkaline phosphatase 10 | 1.64 | 3.32.E-02 |
| CG10116 | uncharacterized protein | 1.57 | 5.23.E-03 |
| **Down-regulated genes** |  |  |  |
| Lcp9 | Larval cuticle protein 9 | 0.089 | 2.43.E-02 |
| CG17107 | uncharacterized protein | 0.121 | 1.74.E-02 |
| Lcp4 | Larval cuticle protein 4 | 0.149 | 4.91.E-04 |
| CG42500 | uncharacterized protein | 0.180 | 4.74.E-03 |
| CG13678 | uncharacterized protein | 0.212 | 2.43.E-02 |
| CG10081 | uncharacterized protein | 0.223 | 3.98.E-06 |
| Cpr47Eb | Cuticular protein 47Eb | 0.321 | 2.07.E-02 |
| CG11737 | uncharacterized protein | 0.399 | 3.72.E-02 |
| CG6277 | uncharacterized protein | 0.423 | 8.20.E-04 |
| bmm | brummer | 0.508 | 6.98.E-04 |
| fax | failed axon connections | 0.547 | 2.22.E-03 |

**Table S2.** Differentially-expressed genes in the adult gut upon larval IMD activation (Fold change>2 or <0.5, adjusted *p*-value<0.05).

| Symbol | Gene Name | Fold Change | *p*-value |
| --- | --- | --- | --- |
| **Up-regulated genes** |  |  |  |
| Mtk | Metchnikowin | 5.95 | 6.63E-04 |
| DptA | Diptericin A | 5.32 | 3.93E-16 |
| CecC | Cecropin C | 4.13 | 9.41E-07 |
| AttC | Attacin-C | 4.10 | 1.60E-02 |
| Dro | Drosocin | 3.80 | 8.27E-06 |
| CecA2 | Cecropin A2 | 3.55 | 6.02E-05 |
| CecA1 | Cecropin A1 | 3.45 | 1.36E-04 |
| lncRNA:CR45045 | long non-coding RNA:CR45045 | 3.38 | 7.33E-04 |
| Listericin | listericin | 2.90 | 8.38E-03 |
| edin | elevated during infection | 2.85 | 6.79E-03 |
| DptB | Diptericin B | 2.72 | 4.67E-03 |
| AttD | Attacin-D | 2.67 | 4.04E-03 |
| CG4269 | uncharacterized protein | 2.62 | 1.42E-02 |
| CG16995 | uncharacterized protein | 2.59 | 1.88E-03 |
| Def | Defensin | 2.26 | 3.17E-05 |
| PGRP-SD | Peptidoglycan recognition protein SD | 2.23 | 7.49E-04 |
| CG10383 | uncharacterized protein | 2.08 | 1.97E-03 |
| Drsl2 | Drosomycin-like 2 | 2.01 | 6.67E-04 |
| Cht4 | Chitinase 4 | 2.01 | 4.87E-02 |
| **Down-regulated genes** |  |  |  |
| CG8745 | uncharacterized protein | 0.303 | 7.33.E-04 |
| CG7567 | uncharacterized protein | 0.311 | 1.29.E-12 |
| CG43673 | uncharacterized protein | 0.331 | 1.60.E-03 |
| Alp10 | Alkaline phosphatase 10 | 0.359 | 1.81.E-04 |
| Alp9 | Alkaline phosphatase 9 | 0.389 | 4.25.E-02 |
| CG3348 | uncharacterized protein | 0.397 | 2.21.E-03 |
| Akh | Adipokinetic hormone | 0.416 | 1.40.E-02 |
| CG32512 | uncharacterized protein | 0.419 | 9.15.E-03 |
| CG7231 | uncharacterized protein | 0.432 | 1.48.E-02 |
| sug | sugarbabe | 0.446 | 9.94.E-05 |
| CG2680 | uncharacterized protein | 0.451 | 7.33.E-04 |
| CG9119 | uncharacterized protein | 0.461 | 3.85.E-04 |
| CG33301 | uncharacterized protein | 0.467 | 4.03.E-06 |
| CG5346 | uncharacterized protein | 0.474 | 3.07.E-07 |
| Ctr1B | Copper transporter 1B | 0.475 | 1.44.E-04 |
| Ser8 | Ser8 | 0.495 | 2.63.E-02 |
| CG15096 | uncharacterized protein | 0.496 | 2.68.E-03 |
| Drat | Death resistor Adh domain containing target | 0.496 | 3.38.E-02 |

**Table S3.**
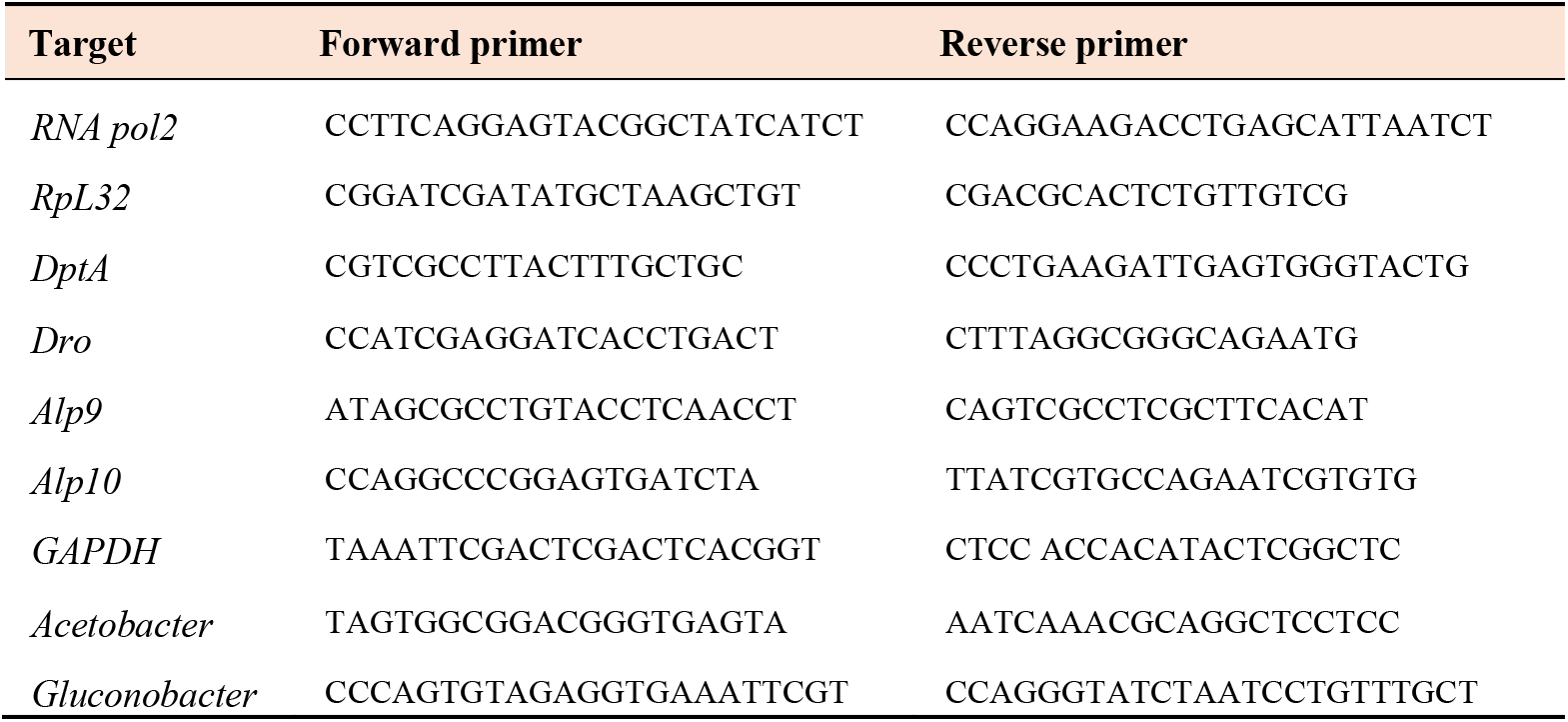
Primer sequences for quantitative PCR analysis.

